# Overstaying in patchy foraging can be explained by behavioral variability

**DOI:** 10.1101/868596

**Authors:** Tyler Cash-Padgett, Benjamin Hayden

## Abstract

Foragers often systematically deviate from rate-maximizing choices in two ways: in accuracy and precision. That is, they both use suboptimal threshold values and show variability in their application of those thresholds. We hypothesized that these biases are related and, more specifically, that foragers’ widely known accuracy bias – over-staying – could be explained, at least in part, by their precision bias. To test this hypothesis, we analyzed choices made by three rhesus macaques in a computerized patch foraging task. Confirming previously observed findings, we find high levels of variability. We then show, through simulations, that this variability changes optimal thresholds, meaning that a forager aware of its own variability should increase its leaving threshold (i.e., over-stay) to increase performance. All subjects showed thresholds that were biased in the predicted direction. These results indicate that over-staying in patches may reflect, in part, an adaptation to behavioral variability.

## INTRODUCTION

Foraging theory provides normative guides for decisions that will maximize long-term reward intake rate in foraging contexts (Charnov 1976; Stephens and Krebs, 1986). Many foragers roughly approximate rate-maximizing behavior (Bixter and Luhmann, 2013; Krebs et al., 1977; McNamara and Houston, 1997; Houston and McNamara, 2014; Kacelnik 1997). Two major apparent deviations from rate-maximizing behavior stand out. ***First***, foragers tend to exhibit much more behavioral variability than they should (Charnov, 1976; Cowie, 1977; Kacelnik, 1984; Lima, 1984; Munger, 1984). This is true even in carefully controlled computerized tasks (Hayden et al., 2011; Blanchard and Hayden, 2014; Eisenreich et al., 2019). A decision-maker that shows variability will harvest less reward than one that does not, because even if their average threshold is correct, it is often incorrect on individual trials. ***Second***, foragers tend to have systematically suboptimal average thresholds and, particularly in patch-leaving contexts, tend to over-stay (Hayden et al., 2011; Blanchard and Hayden, 2015; Cassini et al. 1990; Cassini et al. 1993; Constantino and Daw 2015; Kamil et al. 1988; Kane et al. 2018; Nonacs 2001).

Variability has several potential explanations. One possibility is that there is some factor we are not measuring or considering. These may include internal factors such as deliberate exploration, sensory or motor variability, and cognitive noise (Daw et al. 2006; Evans and Raine 2014; Pyke 1978; Todd and Kacelnik 1993; Ebitz et al., 2019). They may include external factors, such as a limitation in our measurement of foragers’ behavior or our inability to accurately quantify the statistical properties of the environment to which the forager is responding (Brown 1988; Houston and McNamara 1990; Lima and Dill 1990; McNamara and Houston 1987). It can be difficult to disambiguate these possibilities in field studies because much of the information that drives foraging decisions is difficult to measure (Houston and McNamara 2014).

The systematic error in thresholds is more mysterious (Nonacs 2001). Here we propose a new hypothesis, that foragers’ use of suboptimal thresholds may be a rational response to their own variability, and may thus be less costly than it appears. Specifically, if foragers are intrinsically variable and foraging payoff curves are asymmetric (as they typically are), the optimal strategy may be to use a threshold biased in the direction of the shallower slope of the payoff curve. If the forager is both aware of its own variability and unable to reduce it, then the forager may potentially increase its harvest rate by altering its thresholds. Or, more specifically, the forager may be able to mitigate the loss in reward caused by variability by strategically adjusting its threshold.

Here, we examined a dataset consisting of choices made by three foraging macaques in a computerized foraging task (Blanchard and Hayden, 2015). All subjects lived in highly controlled laboratory environments with stable food provisioning and were over-trained on their tasks for months to reduce their subjective uncertainties about task structure as much as possible. All subjects nonetheless showed high behavioral variability, consistent with observations from less controlled field studies. They also showed over-staying. These results suggest that subjects have the flexibility to adjust strategies to account for their own variability.

## METHODS

Three macaque subjects performed a *patch-leaving task* during the collection of data for previously published studies (Blanchard and Hayden 2015; Hayden et al. 2011; Ramakrishnan et al., 2019) and were trained on the task for several months prior. All procedures were approved by the University Committee on Animal Resources at the University of Rochester and were designed and conducted in compliance with the Public Health Service’s Guide for the Care and Use of Animals. Subjects had never previously been exposed to foraging decision-making tasks. Previous training history for these subjects included several types of gambling tasks (Azab and Hayden, 2017 and 2018; Heilbronner and Hayden, 2016; Farashahi et al., 2017) and, in one case, a cognitive set-shifting task (Sleezer et al., 2016).

### Patch-leaving task

Stimuli were colored rectangles a computer screen. A rectangle’s color indicated reward available from that option, and its size represented the delay associated with choosing it. On each trial, subjects acquired fixation on a central point stimulus and were then presented with two targets representing the ‘stay’ in patch and ‘leave’ patch options respectively. After 100 ms, subjects were free to select either target by shifting their gaze to it. Targets shrank in size at a constant rate once selected so that their height provided an unambiguous cue to the delays associated with the two options on every trial. Staying in the current patch always resulted in a 0.6 s “handling time” delay, while the “travel time” associated with leaving varied randomly from patch to patch between 0.5 s and 10.5 s.

If subjects chose to stay in the current patch, a water reward was delivered following the delay. This reward would decrement on each subsequent ‘stay’ decision but would reset upon arrival in a new patch. The initial reward amount for subject C was 200 μL and was decremented by 13 μL for every ‘stay’ decision. The initial reward amounts for subjects E and O were 230 μL and were decremented stochastically, between 12.7 and 15.5 μL per trial, with a mean decrement of 14.1 μL. Further description of the task is available in Hayden et al. (2011).

### Patch-leaving data analysis

We aggregated 14,413 trials across 33 sessions from subject C, 2,010 trials across 4 sessions from subject E, and 1,717 trials across 4 sessions from subject O. In order to control for slight differences in travel times and handling times between the two datasets, we analyzed the subjects’ foraging behavior in terms of the number of ‘stay’ decisions per patch. Mean patch residence time was defined as the mean number of trials spent in-patch across all patches. Reward rate was defined as the mean amount of water received per patch across all trials.

### Patch-leaving task simulation

To calculate the impact of noise on a subject’s optimal threshold value, we constructed a simulation of the patch-leaving task as described above. Each simulated subject ran 10,000 behavioral sessions of 439 trials (the average number of trials per session over all three datasets). Patch-leaving thresholds ranged from 1 to the maximum number of rewarded trials depending on initial reward and decrement size (18 for subject C, 17 for subjects E & O). We uniformly distributed the possible thresholds across the patches in each session. The expected reward rate for a given threshold (Figure 2B) was calculated as the average reward rate from all patches, in all sessions, where that threshold was in effect.

**Figure 1.**
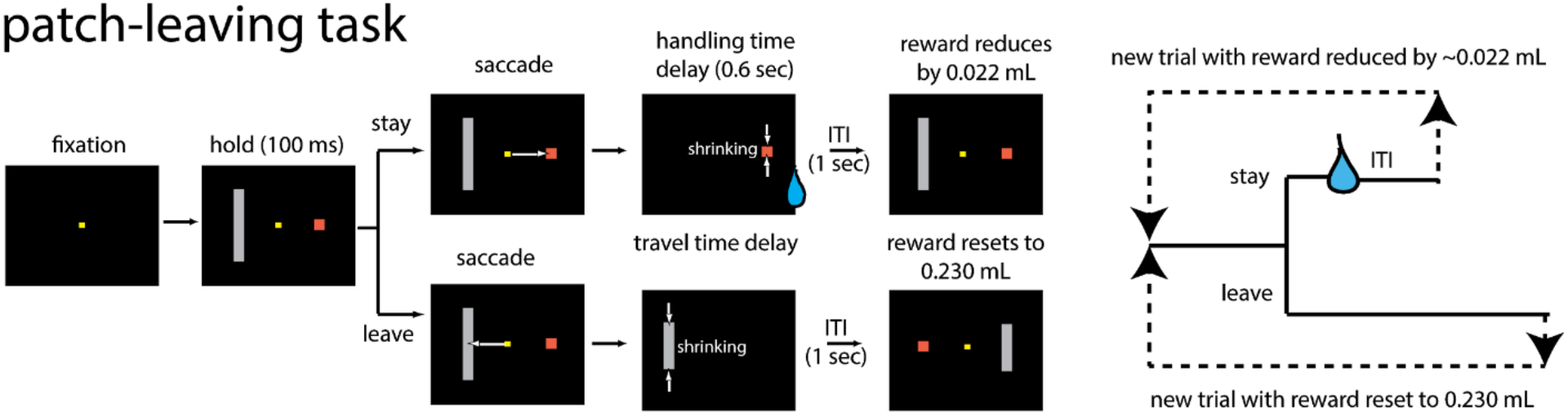
*Task*. After fixation, the subject views two targets, and chooses one by shifting gaze to it. Choice of the orange rectangle (stay in patch) yields a short delay and reward whose value diminishes by 13 μL/trial. Choice of the gray rectangle (leave patch) yields no reward and a long delay (travel time) whose duration is indicated by the height of the bar, and resets the value of the orange rectangle at 230 μL. Patch travel time varies randomly from patch to patch between 0.5 to 10.5 s.

**Figure 2.**
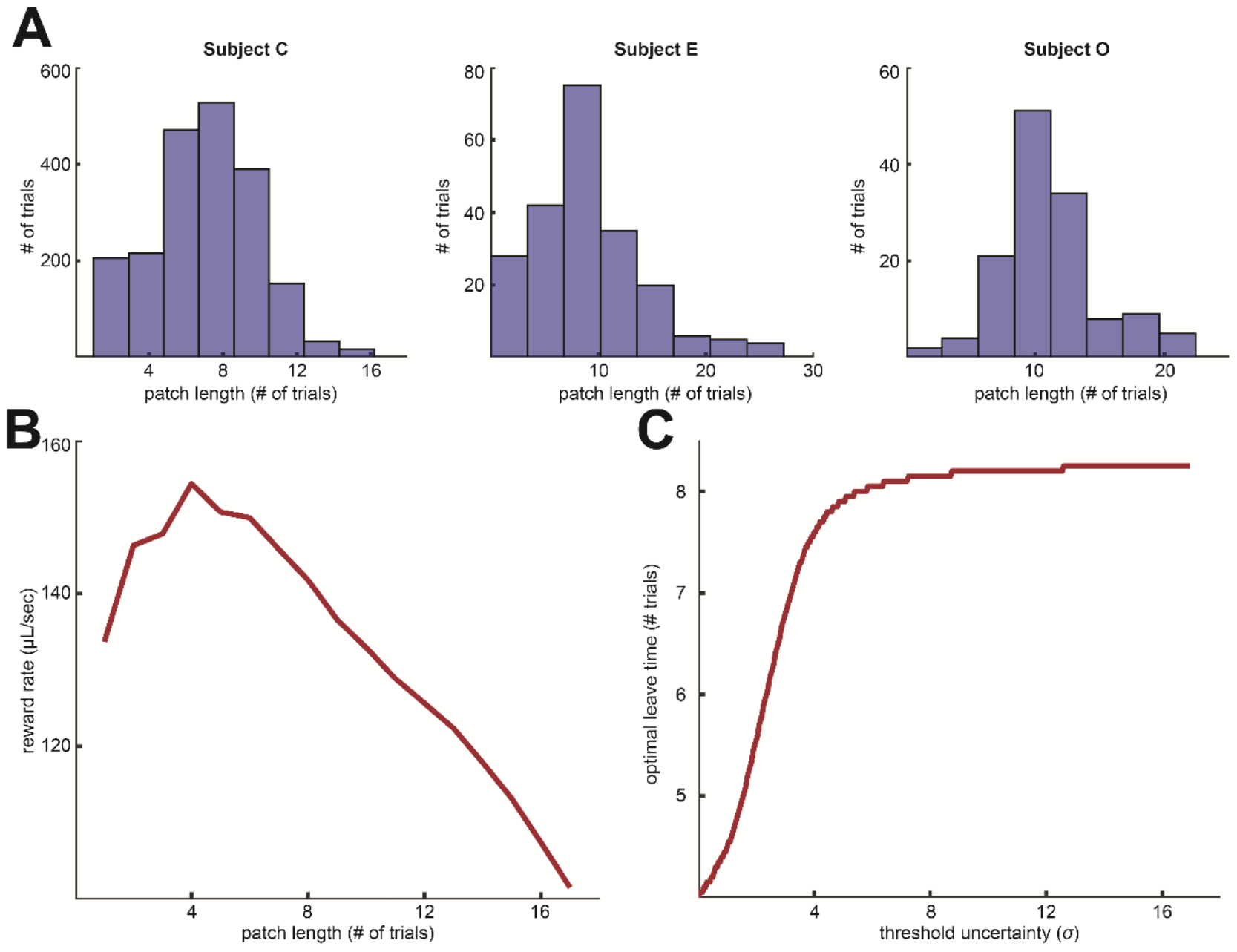
*Patch-leaving behavior and simulation*. **(A)** Patch length distributions for each subject denoted by number of trials (‘stay’ decisions). **(B)** The reward rate available via behavioral policies of leaving after a threshold number of trials, per simulation. **(C)** The optimal patch leaving threshold as a function of a set amount of Gaussian noise applied to the threshold (quantified by the standard deviation *σ*).

The total accumulated reward in a given patch was based on the number of ‘stay’ decisions, initial reward, and reward decrement amount. We therefore normalized the reward rate to the number of trials, rather than raw time, spent in-patch. Travel time was normalized by dividing, for each patch residence duration, the mean and standard deviation of travel times observed during behavioral testing, respectively, by total patch residence time. Travel time for each simulated patch was then drawn from a normal distribution based on the empirically observed mean and standard deviation for that particular normalized patch residence duration. Since we lacked precise timing information for subject C, this normalized travel time function was calculated based on behavioral data from subjects E and O.

To simulate the effect of threshold noise on its optimal value, we interpolated the reward curve twenty-fold and performed trapezoidal numerical integration for a Gaussian-distributed matrix centered on each candidate threshold value. For standard deviations ranging from zero to one half of the maximum number of rewarded trials, we calculated the noise-adjusted threshold value that maximized the area under the reward rate curve.

## RESULTS

Three macaques (subjects C, E, and O) performed a series of trials in which they chose between remaining in a patch or leaving it. Remaining led to a short “handling time” (delay of 600 ms) and leaving led to a longer “travel time” (delay of 0.5 to 10.5 seconds, see **Methods** for details). Travel time varied randomly between patches and was signaled unambiguously within each patch. To control for slightly differing travel and handling time regimens between our three subjects, we defined patch residence time as the number of decisions to stay in a patch.

Both subjects showed residence times that were close to, but longer than, the dictates of foraging theory (Stephens and Krebs, 1986), which yields an optimal patch residence time of **4.01 (± 0.74) trials** for subject C’s version of the task and **4.16 (± 0.78) trials** with subject E and O’s slightly different version of the task. In practice, all three subjects’ mean patch residence times were significantly longer than optimal: **5.84 (± 0.08) trials** for subject C, **9.26 (± 0.41) trials** for subject E, and **12.09 (± 0.42) trials** for subject O (all significant, two-sided Student’s t-test, p < 0.0001)

Subjects’ trial-to-trial behavior in this task showed significant variability, which was costly. A forager with no variability would have obtained a reward rate of **168.4 μL/patch** with subject C’s observed threshold. Subject C’s actual reward rate, however, was **128.4 μL/patch**. Similarly, a forager with no variability would have obtained a reward rate of **174.4 μL/patch** with subject E’s observed threshold, but the subject’s actual reward rate was **161.0 μL/patch**. Finally, while a forager with no variability would have obtained a reward rate of **147.1 μL/patch** with subject O’s observed threshold, the subject’s actual reward rate was **135.1 μL/patch**.

We calculated the trial-to-trial variability in each subject’s patch leaving behavior as the standard deviation in its threshold over a large sample of bootstrapped 500-trial bins. Based on an empirical threshold standard deviation of **4.15 trials/patch**, Subject C’s uncertainty-adjusted optimal patch residence threshold was **7.40 trials**. In other words, optimizing given his variability can account for some of his observed rate adjustment (**+1.84 trials**) relative to optimal, assuming no variability (**+3.40 trials**). A similar pattern held for subjects E and O, with respective standard deviations of **6.02 trials per patch** and **5.04 trials per patch** both suggesting a compensatory increase in threshold. While their behavioral adjustments are in the same direction as suggested by accounting for uncertainty, however, the observed adjustments are actually larger than optimal (**+4.26** trials/patch observed vs. **+3.05** predicted for subject E, and **+7.09 trials** observed vs. **+2.95 trials** predicted for subject O).

## DISCUSSION

We examined previously collected datasets of rhesus macaques performing a computerized patch foraging task (Blanchard and Hayden, 2015). Macaques showed two behavioral patterns that are characteristic of many foragers. First, they showed a systematic deviation from an optimal foraging threshold by over-staying. Second, they showed a strong and costly variability in behavior. We propose that these two phenomena are at least partly related. First, we conjecture that behavioral variability is for some reason unavoidable (and our results do not offer any explanation for it). Given this unavoidability, we show that the reward-maximizing threshold increases. We then show that all three subjects showed changes in the direction that yielded improved harvest rates.

Previous studies of foraging behavior have suffered two limitations that ours avoids. First, many studies are limited by data quantity. By using well-trained macaques performing a computerized task, we were able to analyze several thousand trials. Second, many studies have sources of unmeasurable noise. These include difficulty quantifying the details of the environment and difficulty knowing that the forager has had sufficient experience with the environment to make the same calculations we as observers would. The carefully controlled nature of our computerized task enables us to fix all relevant task variables and ensure overtraining on those specific tasks. We were further able to reduce unmeasured variability by controlling animals’ learning environment in their juvenile and adult lives and by ensuring a relatively stable food supply over that entire period of time.

It is perhaps surprising that subjects’ behavioral variability on this task was so high despite the stable environment and months of preparatory task training. This result replicates a classic finding from a seminal study (Charnov, 1976). Our own study does not shed any light on the source of behavioral variability. A long tradition in foraging theory emphasizes the importance of “informational constraint” on suboptimal behaviors (Eisenreich et al. 2017; Fernández-Juricic et al. 2004; Pearson et al. 2010; Stephens 2002). Other factors may also be relevant, including information-seeking (Blanchard et al. 2015; Bromberg-Martin and Hikosaka 2009), temporal discounting (Kane et al. 2018), curiosity (Wang and Hayden 2019), computational noise, or exploratory behavior (Ebitz et al. 2018; Pearson et al. 2009; Wilson et al. 2014). Whatever the source, our results highlight an intriguing fact about monkeys’ variability: they have adjusted their behavior as if they have learned to reduce its costs. This suggests that they may be unable or unwilling to directly change it, but flexible enough to adjust surrounding aspects of their behavior.

Foraging is a major driver of the animal brain (Calhoun and Hayden 2015; Hayden 2018; Murray and Rudebeck 2013; Passingham and Wise 2012; Pearson et al. 2014). As such, a science of the neural basis of choice ought to go hand in hand with an understanding of foraging psychology. Systematic deviations from optimal foraging provide an important measure of the forces of evolution; because they are costly, they likely reflect the existence of tradeoffs that are not obvious to evolutionary biologists. Many of these likely come from constraints on the costs of computing foraging strategies in full.

## AUTHORS’ CONTRIBUTIONS

TCP performed all data analysis, designed the simulations, and drafted the manuscript. BYH posited the study’s hypothesis and critically revised the manuscript. Both authors gave final approval for publication and agree to be held accountable for the work performed therein.

## Acknowledgements

This work is supported by a CAREER award from NSF (BCS1253576) and a R01 from NIH (DA038615) to BYH. We thank Meghan Castagno, Marc Mancarella, Caleb Strait and Tommy Blanchard for assistance with data collection, and the rest of the Hayden lab for valuable discussions.

